# *E. coli* Leucine-responsive regulatory protein bridges DNA *in vivo* and tunably dissociates in the presence of exogenous leucine

**DOI:** 10.1101/2022.03.07.483379

**Authors:** Christine A. Ziegler, Peter L. Freddolino

## Abstract

Feast-Famine Response Proteins are a widely conserved class of global regulators in prokaryotes, the most highly studied of which is the *E. coli* leucine-responsive regulatory protein (Lrp). Lrp senses environmental nutrition status and subsequently regulates up to one-third of the genes in *E. coli*, either directly or indirectly. Lrp exists predominantly as octamers and hexadecamers (16mers), where leucine is believed to shift the equilibrium towards the octameric state. In this study, we analyzed the effects of three oligomerization state mutants of Lrp in terms of their ability to bind to DNA and regulate gene expression in response to exogenous leucine. We find that oligomerization beyond dimers is required for Lrp’s regulatory activity, and that contrary to prior speculation, exogenous leucine modulates Lrp activity at its target promoters exclusively by inhibiting Lrp binding to DNA. We also find evidence that Lrp binding bridges DNA over length scales of multiple kilobases, revealing a new range of mechanisms for Lrp-mediated transcriptional regulation.

## Introduction

Bacterial transcription is controlled in a hierarchical manner, in which a small number of highly expressed global regulators exert control over the expression of hundreds of transcription factors and downstream target genes [1,2]. *E. coli* has seven such global regulators: CRP, IHF, FNR, Fis, ArcA, H-NS, and Lrp [1]. Of these global regulators, Lrp (<u>L</u>eucine-responsive <u>R</u>egulatory <u>P</u>rotein) is of particular interest due to its crucial role in nutrient sensing and its high conservation among prokaryotes; nearly half of sequenced bacteria and almost all sequenced archaea contain one or more Lrp homologs [3]. While Lrp was originally identified in 1973 by the Oxender Lab [4] and has since been shown to regulate as many as 32% of *E. coli* genes either directly or indirectly [5], little is known about the precise mechanism by which Lrp regulates its target genes [6].

Lrp is a small (18.8kDa), basic (pI=9.2), and highly abundant DNA-binding protein (∼3,000 dimers/cell) in *E. coli [7]*, and is thought to sense environmental nutritional status via exogenous L-leucine levels and to regulate gene expression accordingly [7]. The *E. coli* Lrp regulon consists of genes involved in amino acid metabolism, one-carbon metabolism, peptide and amino acid transport, flagella and fimbriae biosynthesis, osmotic stress, acid stress, and other virulence factors [5,8–10], and Lrp is known to regulate 70% of the genes involved in the transition from logarithmic growth to stationary phase as nutrients are depleted [11]. To accomplish its regulatory activities, Lrp was shown to utilize six different modes of gene regulation: Lrp can activate or repress its target genes, and for each target leucine can have no effect, augment Lrp’s effect, or inhibit Lrp function [8,9]. Through several *in vitro* studies, the Calvo Lab hypothesized that Lrp exists predominantly as a hexadecamer (16mer) when L-leucine concentrations are low (such as in minimal media), but that the oligomeric state switches to predominantly Lrp octamers in the presence of L-leucine, suggesting that leucine modulates Lrp function by altering Lrp’s oligomeric state [12–14]. However, this hypothesis does not fully explain how leucine can augment Lrp function at some promoters, have no effect at others, and somehow inhibit the rest.

In this study, we determined a more precise mechanism by which exogenous L-leucine modulates Lrp gene regulation by analyzing the global effects of leucine supplementation in the media on Lrp binding to DNA and its subsequent effects on gene expression via paired Lrp ChIP-seq and RNA polymerase ChIP-seq. In order to further investigate how leucine supplementation and oligomeric state affect Lrp gene regulation, we also analyzed the effects of exogenous leucine on a “dimer-only” Lrp mutant (ΔC11), as well as two “octamer-only” Lrp mutants, D114E and L136R, that were previously identified by Platko and Calvo [12,13,15]. Through these high-throughput experiments and targeted follow-up case studies, we determined that exogenous L-leucine modulates Lrp gene regulation by inhibiting Lrp binding to DNA at its target promoters. Lrp likely exists in an equilibrium of various higher-order oligomeric complexes that bridge DNA in minimal media depending on the DNA sequence, and only the strongest Lrp binding sites maintain Lrp bound to the DNA in the presence of L-leucine. The dimer-only mutant is unable to bind DNA under any condition tested, demonstrating that Lrp requires oligomers larger than a dimer in order to bind DNA. The Lrp L136R mutant, which we show behaves like WT Lrp in minimal media, is unable to sense exogenous leucine and therefore maintains higher-order oligomeric nucleoprotein complexes on target promoters, even in the presence of leucine. Meanwhile, the D114E mutant binds to the canonical Lrp sites as well as additional sites next to known binding sites in minimal media, suggesting that it stabilizes higher-order Lrp oligomers that bridge DNA. Thus, while neither the L136R nor D114E mutants are strictly “octamer only”, the differing effects of these mutations on Lrp’s DNA binding and regulatory output allow us to parse out the effects of ligand binding and oligomerization.

## Results

### Construction of a faithful genetic background for analyzing Lrp function

To the best of our knowledge, many previous studies of *E. coli* Lrp utilized strain backgrounds with mutations in one or more of the three acetohydroxyacid synthase (AHAS) genes: *ilvBN, ilvGM*, or *ilvIH*, which are involved in the regulation and biosynthesis of branched-chain amino acids [16]; for example, the commonly used K12 strain MG1655 has a frameshift mutation in *ilvG*, whereas *ilvB* and *ilvIH* were mutated in some of the original lrp studies. Because Lrp responds strongly to the presence of L-leucine and regulates branched-chain amino acid transport and biosynthesis, we opted to use the RL3000 lab strain of *E. coli* for all of our experiments, which is a derivative of MG1655 harboring a functional *ilvG* gene, and therefore contains all three fully functional AHAS isozymes [17].

To preserve Lrp regulation and copy number, we chose to integrate all Lrp mutants directly into the genome; however, the native *lrp* locus was difficult to modify without disrupting several LexA sites downstream of *lrp* or altering *ftsK* expression, which is an essential gene involved in cell division that is located immediately downstream of *lrp*. To circumvent these issues, we instead deleted the entire open reading frame of *lrp* at its native locus, leaving behind an FRT scar (*lrp::scar*), and integrated a cassette containing the native *lrp* promoter and WT *lrp* gene flanked by strong bidirectional terminators into a locus immediately downstream of the native *thyA* gene (Fig. 1A). We chose the *thyA* locus because *thyA* encodes Thymidylate Synthase, which can be used as a Lrp cassette-linked selectable marker on media lacking thymidine, and because the *thyA* and *lrp* loci are roughly equidistant from the origin and therefore have similar DNA copy numbers. To investigate the effect of oligomeric Lrp mutants on gene expression, we also integrated the “dimer-only” ΔC11 and the “octamer-only” D114E and L136R *lrp* genes into the thyA locus using the same approach (Table S1).

**Figure 1:**
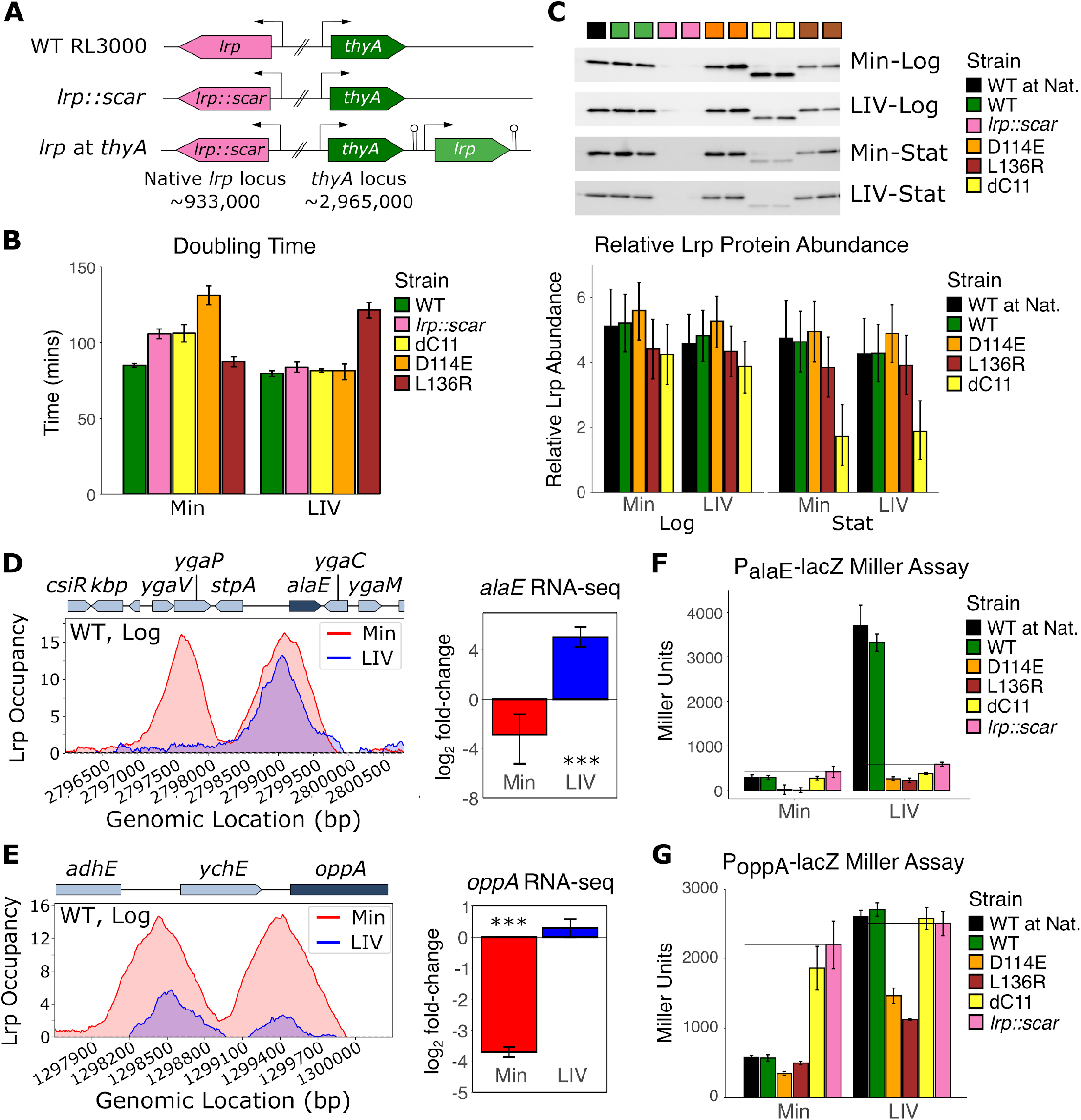
Lrp mutants display unique and consistent phenotypes. **A**. WT RL3000 contains *lrp* at the native locus (pink), whereas in the *lrp::scar* and *lrp* at *thyA* strains the *lrp* ORF at the native locus has been fully removed. The *thyA* locus in these strains contains either *thyA* alone (*lrp::scar*) or *thyA* immediately upstream of a bidirectional-terminator-flanked *lrp* gene with its native promoter (*lrp* at *thyA*). Arrows represent promoters and hairpins represent bidirectional terminators. **B**. Doubling times of *lrp* strains in Min and LIV M9 minimal media were calculated during mid-log phase and each bar represents the mean of two biological replicates each of two independently-constructed lineages of each strain. Error bars represent the standard deviation. All *lrp* variants in this experiment are at the *thyA* locus, including WT. **C**. Top, a representative Western Blot for each condition+time point showing both lineages of each strain. Bottom, inferred Lrp abundances under each condition obtained by fitting a Bayesian model for Lrp protein levels (based on pixel density of Western Blots divided by the sum of pixel density from a total protein blot) across strains, conditions, and both biological and lineage replicates. Error bars indicate a 95% credible interval. For model fitting, *lrp::scar* was excluded because there was no Lrp present in the Western Blots for these samples. **D**. Left, re-plotted WT Lrp ChIP-seq data from [5] for the *stpA-alaE* region during mid-log phase, where the y-axis represents the *lrp::scar*-subtracted z-scores of Lrp-ChIP relative to input for Min media (red) and LIV media (blue). Right, re-plotted log_2_-fold change of *alaE* transcript levels from [5] of WT Lrp cells relative to *lrp::scar* cells during mid-log phase in Min media (red) and LIV media (blue). Error bars represent a 95% confidence interval. **E**. Same as D), except for the *oppA* region. **F**. Bars represent the mean Miller Units from the P_alaE_-*lacZ* reporter Miller Assay of two technical replicates from each of two lineage replicates for each strain in each condition (Min and LIV) at mid-log phase. Error bars represent the standard deviation. Black horizontal lines represent the Miller Units for the *lrp::scar* strain in that condition. **G**. Same as F), except with the *P*_*oppA*_*-lacZ* reporter construct.

### Oligomeric Lrp mutants show altered growth phenotypes and regulatory outputs

With a safe system for genetically modifying *lrp* in hand, we proceeded to investigate how WT Lrp as well as various Lrp oligomeric mutants (D114E, L136R, and ΔC11) affected the general physiology of *E. coli* cells in minimal media (Min) and the same media supplemented with the branched-chain amino acids (LIV). To our surprise, the strain harboring the “dimer-only” ΔC11 had a similar doubling time as *lrp::scar* in media with and without leucine (Fig. 1B). Intriguingly, while D114E and L136R were both originally identified as “octamer-only” mutants and were therefore expected to behave similarly, D114E had a strong growth defect in Min media that was restored in LIV media, whereas L136R had the opposite effect on growth rate in Min and LIV media (Fig. 1B). We next investigated the Lrp protein levels in each of our strains via Western blotting and confirmed that WT Lrp expressed from the *thyA* locus was present at comparable levels as WT Lrp expressed from the native *lrp* locus (Fig. 1C and Table S2). Despite ΔC11 strains growing at comparable rates to *lrp::scar*, the dimer-only ΔC11 protein is expressed at only slightly lower levels than WT Lrp during logarithmic growth (>97% posterior probability of being less than the WT level based on our Bayesian analysis; here we abbreviate the posterior probability of a difference in the observed direction as P_diff_); however ΔC11 levels are substantially reduced over the course of the growth curve (P_diff_ >99%). We obtained weaker evidence that D114E Lrp was more highly expressed than WT Lrp in all conditions tested (P_diff_>76%), while L136R Lrp was expressed at lower levels than WT Lrp (P_diff_>80%) (Fig. 1C and Table S2), which may partially explain the differential growth rates of these two octamer-only mutants; however it should be noted that *lrp* is autoregulated. Thus, contrary to previous expectations regarding the octamer-only D114E and L136R Lrp variants, they in fact appear to have distinct responses to changing nutrient conditions.

To assess how these oligomeric Lrp mutants affect gene expression at well-studied leucine-Lrp regulated promoters, we next constructed *lacZ* reporter strains at the native *alaE* and *oppA* promoters. *alaE* and *oppA* were selected due to their unique Lrp binding patterns and response to leucine observed by Kroner *et al*. [5]. The *alaE* promoter contains two strong Lrp peaks in Mn media that cause a slight repression of *alaE*, whereas only one of the two Lrp peaks is maintained in LIV media, causing a strong activation of *alaE* (Fig. 1D). On the other hand, WT Lrp forms two strong peaks at the *oppA* promoter in minimal media that repress *oppA*, but these Lrp peaks are reduced to almost no Lrp binding in LIV media, and repression of *oppA* becomes de-repressed (Fig. 1E). Our WT and *lrp::scar* Miller assays reproduced the findings from Kroner *et al;* Lrp slightly represses *alaE* in minimal media, but strongly activates *alaE* expression in LIV (Fig. 1F), whereas Lrp binding represses *oppA* in minimal media, which is relieved in LIV (Fig. 1G). Additionally, the regulation at both of these promoters by WT Lrp expressed from the native locus (black bars) and WT Lrp expressed from the *thyA* locus (green bars) are comparable across conditions, validating that our *lrp-thyA* constructs mimic WT strains. As was observed in the growth rate experiment, dimer-only Lrp (ΔC11) behaved like *lrp::scar* at *oppA* (Fig. 1G), but appeared to have a slight repressive effect on *alaE* relative to *lrp::scar* regardless of whether LIV was in the media (Fig. 1F), suggesting that Lrp ΔC11 has a potential regulatory effect for at least some Lrp-regulated promoters.

Both D114E and L136R Lrp repressed *oppA* in Min media and maintained most of their repression in LIV (Fig. 1G). These two octamer-only Lrp mutants also behaved similarly at the *alaE* promoter, where they both repressed *alaE* even more strongly than WT Lrp in minimal media, and this repression was relieved in LIV (Fig. 1F). Unlike WT Lrp, for both octamer-only mutants, *alaE* expression was not activated by LIV. Collectively, these data confirm that our genetic system reproduces previous data and also demonstrate a potential regulatory role of dimer-only ΔC11. However, at least for the two target promoters considered here, the octamer-only D114E and L136R variants have similar regulatory effects, which cannot explain the opposite growth phenotypes of these two strains. To address the discrepancy between observed regulation and growth phenotypes, we next took a high-throughput approach to globally assess Lrp binding and regulation of gene expression in response to different Lrp mutants and conditions.

### Paired Lrp- and RNAP-ChIP seq reproduce previous findings and reveal unique binding patterns for each of the oligomeric Lrp mutants

In order to obtain genome-wide information on the interplay of *lrp* oligomeric state and physiological conditions with Lrp’s binding state and regulatory output, we performed a series of ChIP-seq experiments targeting both Lrp and RNA polymerase (RNAP). WT Lrp, *lrp::scar*, D114E, L136R, and ΔC11 (Table S1) strains were grown in glycerol-ammonia M9 minimal media (Min) or glycerol-ammonia M9 minimal media supplemented with all three branched-chain amino acids (LIV) and harvested for ChIP-seq at mid-log phase and stationary phase, as shown in Figure 2A. To ensure reproducibility, each strain was constructed twice (lineage A or B), and two biological replicates were performed on different days for each strain in each lineage for each condition. After Illumina sequencing and read processing, we called all significant Lrp peaks within each sample and merged these peaks into one list of all detectable Lrp binding regions (see Methods for details). In total, we detected 793 Lrp binding regions across the genome. Figure 2B is a row-normalized heatmap of Lrp occupancy (color) at each Lrp binding site (Y-axis) for each sample (X-axis), where both the rows and columns are clustered by similarity.

**Figure 2:**
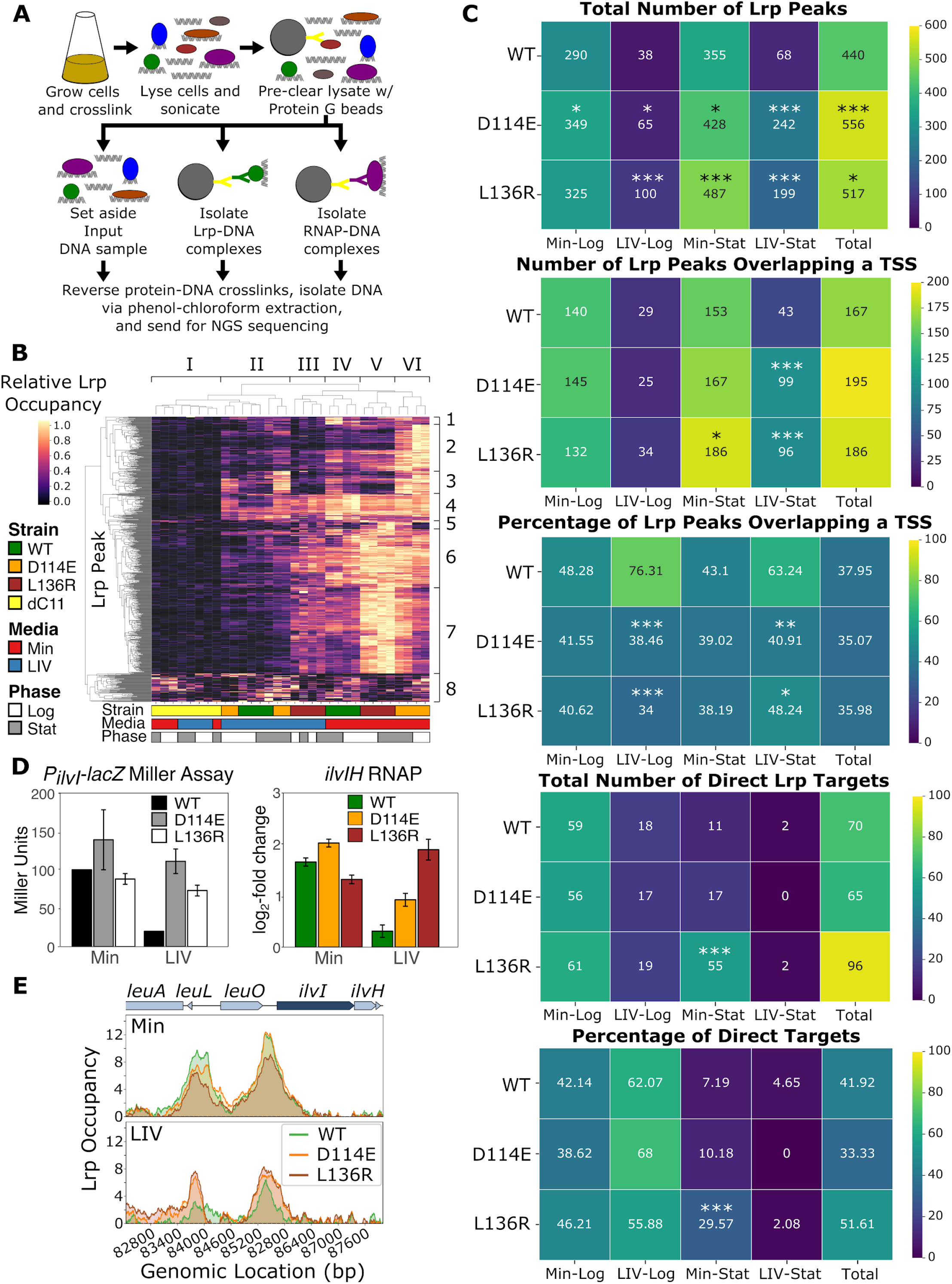
Paired Lrp- and RNAP-ChIP-seq reveals several key features of Lrp binding and regulatory effects. **A**. Schematic of paired Lrp- and RNAP-ChIP-seq experimental workflow, where proteins are represented as colored circles, the magnetic Protein G beads are represented as large gray circles with a Y-shaped yellow antibody attached to the surface, Lrp and RNAP are represented by green and purple circles, respectively, whereas the Lrp and RNAP-specific antibodies are represented by green and purple Y-shapes, respectively. See Materials and Methods for a detailed protocol. **B**. Heatmap of Lrp occupancy at all Lrp binding regions across all strains, media, and timepoints, where the x-axis contains each sample, the y-axis represents each of the 793 significant unique Lrp binding sites that were identified in at least one strain in at least one condition, and the color at each position represents the row-normalized Lrp occupancy in that sample at that location (Lrp occupancy divided by the maximum value in each row). Each column represents the mean of two biological replicates, and each lineage replicate is plotted separately. Both the x- and y-axes were clustered by Euclidean distance (taking the average distances between cluster members), generating Groups I-VI (samples) on the x-axis and Classes 1-8 (binding site categories) on the y-axis. **C**. Heatmaps of total Lrp peaks, Lrp peaks at promoters, and direct Lrp targets for each strain and condition. For each plot, the numbers in each box represent the total number (or percentage) of unique Lrp peaks or transcriptional units (TUs) across both biological replicates of both lineage replicates for each strain in each condition. The “Total” column in each heatmap represents the total number of unique Lrp peaks or TUs within each strain across all conditions and is not simply the sum across the row. Stars represent the significance relative to WT Lrp in that condition, where * indicates p<0.05, ** indicates p<0.01, and *** indicates p<0.001. Significance for total numbers was calculated by comparing rates using the `poisson.test` function in R, whereas significance of percentages was determined using a 2-sample test for equality of proportions using the `prop.test` function in R. **D**. Comparison of re-plotted P_ilvIH_-*lacZ* Miller assays by Platko and Calvo [15] for the Lrp variants shown (left) with our RNAP-ChIP data for *ilvIH* (right). The RNAP-ChIP data shows the log_2_-fold change in *lrp::scar*-subtracted RNAP occupancy relative to Input for *ilvIH* for WT, D114E, and L136R Lrp at mid-log phase in Min and LIV media. Each bar represents the mean of two biological replicates for each of the two lineages, and the error bars represent the standard error. **E**. The y-axis represents the mean Lrp occupancy (*lrp::scar*-subtracted rz-log-ratio of Lrp relative to Input) across both biological and lineage replicates for each strain in Min (top) and LIV (bottom) during mid-log phase at the *ilvIH* locus. Changes in RNAP occupancy in D) are roughly correlated with Lrp occupancy upstream of *ilvIH*.

The samples (X-axis of Fig. 2B) are clustered into six groups (I-VI). Group I consists of dimer-only Lrp ΔC11 samples and generally exhibits low binding to DNA at most Lrp binding sites, indicating that this dimer-only mutant is deficient in DNA binding. For this reason, we exclude Lrp ΔC11 from further ChIP-seq analysis. Group II is a mixture of WT and D114E samples both grown in LIV, which also exhibit lower Lrp binding, with the exception of a few highly occupied sites. Groups III (L136R-LIV) and IV (WT-Min) cluster closely together, indicating that at the genome-wide level L136R often fails to recognize exogenous leucine, and shows a binding profile in the presence of leucine similar to that of WT Lrp in minimal media. Similarly, Groups V (L136R-Min) and VI (D114E-Min) cluster closely together and exhibit the highest Lrp binding of all samples. Given that D114E and L136R were both identified as “octamer-only” mutants, it was surprising that D114E exhibits exogenous leucine sensitivity (like WT Lrp), but L136R is deficient in leucine-sensing. Furthermore, we were surprised to see such strong binding of both octamer-only mutants to DNA in Min media. These changes in Lrp binding between D114E and L136R across conditions could explain the differences in growth rates between them (shown in Fig. 1B). D114E likely has a slower growth rate in Min media due to the presence of many novel Lrp binding sites under this condition, which likely cause suboptimal regulation of pathways important for growth. Conversely, L136R may exhibit its growth defect in LIV media since this strain exhibits more Lrp binding in LIV than any other strain (comparable to WT Lrp binding in Min) and essentially acts as if it were growing in minimal media, despite the presence of leucine and other branched chain amino acids.

It is also useful to consider the clustering of Lrp binding regions (Y-axis of Fig. 2B), which separate into eight distinct classes (1-8). Class 1 consists of WT and D114E Min-Stat specific sites. Class 2 contains the D114E-Min specific binding sites, whereas Class 3 consists of regions bound by D114E regardless of the presence of leucine, demonstrating D114E’s leucine insensitivity at only a subset of sites. Class 4 is a broad class of sites that are relatively leucine insensitively bound by WT, D114E, and L136R; these are likely some of the strongest Lrp binding sites since they remain occupied by Lrp even in LIV. Class 5 contains L136R-Stat specific sites, of which there are only a few. Conversely, the large Class 6 contains WT, D114E, and L136R Min specific peaks, as well as the L136R LIV peaks, which thus represent the ‘normal’ targets of Lrp in the minimal media condition and further demonstrate that L136R in LIV acts like WT in Min. Class 7 consists of predominantly L136R-Min and D114E-Min-Log peaks. Finally, Class 8 is made up of non-specific Lrp binding and could be attributed to noise in our peak-calling method. Overall, the general trend is that Lrp is more broadly bound to DNA in minimal media than in leucine-containing media (with the exception of L136R, which cannot sense leucine). Furthermore, there are many novel sites occupied by D114E and, to a much lesser extent, L136R. Thus, a genome-wide perspective reveals substantive differences in behavior of the two octamer-only (D114E and L136R) mutants, with D114E showing a fundamentally altered profile of binding locations whereas L136R binds more strongly to canonical Lrp sites in the genome and persists in binding regardless of the presence of leucine.

To quantify differences in Lrp binding across genotype and condition, we first calculated the number of significant peaks, both for each condition and overall within a genotype (Fig. 2C). Overall, WT Lrp had the fewest peaks (440 overall), with L136R and D114E having significantly more peaks (517 and 556, respectively). However, the overall distribution of peaks within a genotype across conditions remained similar, with Min-Log and Min-Stat having the most peaks and LIV-Log having the least. We next wanted to investigate the role of these Lrp peaks in gene regulation and saw, perhaps unsurprisingly, that Lrp binding sites are strongly enriched at promoters (Fig. 2C), with 37.95% of WT, 35.07% D114E, and 35.98% of L136R peaks overlapping at least one annotated transcription start site (TSS), representing significant enrichments over what would be expected by chance (p<0.01, approximate permutation tests). Of the Lrp peaks overlapping a TSS, we next calculated the number of transcriptional units (TUs) that are directly regulated by Lrp (i.e. Lrp binds to the TSS and there is a significant Lrp-dependent change in RNAP occupancy), and found that 70/167 (41.92%) of WT-, 65/195 (33.33%) of D114E-, and 96/186 (51.61%) of L136R-regulated TUs were under direct Lrp regulation in at least one condition tested. Given that Lrp is strongly enriched at promoters, it is likely that the remaining poised Lrp peaks could become direct regulators of gene expression under a condition not tested in this experiment, as was previously suggested [5]. The role of Lrp sites within open reading frames remains unclear, although, given that the presence or absence of Lrp at these sites does not significantly affect the respective gene expression, it is likely that these sites serve as decoys in order to correctly titrate Lrp levels at functional sites [18,19].

To further validate our system and confirm that the high-throughput paired Lrp- and RNAP-ChIP-seq data reflected that of previous smaller-scale Lrp findings, we examined the regulation of the *ilvIH* locus. In their 1993 study, Platko and Calvo identified several Lrp mutants (including D114E and L136R) and conducted Miller assays on *ilvIH-lacZ* fusion strains grown in Min or LIV media to assess Lrp function at this promoter [15]. In general, they found that WT Lrp (when compared with a *Δlrp* strain) strongly activated *ilvIH* in Min, but not in LIV media, whereas the D114E and L136R strongly activated *ilvIH* in both Min and LIV media (re-plotted in Fig. 2D, left). The change in average RNAP occupancy (relative to *lrp::scar*) across the *ilvIH* transcript in our experiments (Fig. 2D, right) revealed the same patterns of Lrp regulation, demonstrating strong reproducibility between our experiments, despite being performed decades apart in different background strains, different laboratories, with a different method, and in slightly different media compositions. Furthermore, we noticed a striking correlation between the degree of Lrp binding at the *ilvIH* promoter and *ilvIH* expression across strains and media, where leucine seemed to reduce WT Lrp binding to the promoter, but not D114E or L136R (Fig. 2E).

### Leucine inhibits WT Lrp binding to DNA

One of the major outstanding questions regarding *E. coli* Lrp is how leucine can have no effect on Lrp at some promoters, have a concerted effect with Lrp on others, and have an inhibitory effect on Lrp for the remainder (proposed in [8] and [9]). To globally investigate the roles of leucine and growth phase on WT Lrp binding to DNA, we called all significant WT Lrp occupancy peaks, totalling 440 across all WT Lrp conditions, and plotted Lrp occupancy within each sample at each of these sites (Fig. 3A). It is immediately apparent that leucine reduces or completely prevents Lrp binding at nearly all of these regions, suggesting that exogenous leucine ultimately acts to inhibit Lrp binding to DNA. To investigate the relationship between WT Lrp binding at promoters and the changes in expression at the accompanying transcriptional units (TUs), we next plotted the occupancy traces of Lrp within a 2kb window of all 142 annotated TSSs overlapped by a WT Lrp peak (Fig. 3B) alongside the Lrp-dependent change in RNAP occupancy at the associated TU (here we use the presence of RNA polymerase in the gene body as a proxy for active transcription, and quantify it accordingly); a positive value indicates Lrp-mediated activation and a negative value denotes Lrp-mediated repression (Fig. 3C). In accordance with the trend from Figure 3A where exogenous leucine reduces Lrp occupancy at Lrp binding sites, the LIV-Log condition in Figure 3B has a greatly reduced Lrp occupancy at TSSs relative to the Min-Log condition, and these changes in Lrp occupancy at the TSSs are associated with substantial changes in gene expression (Fig. 3C).

**Figure 3:**
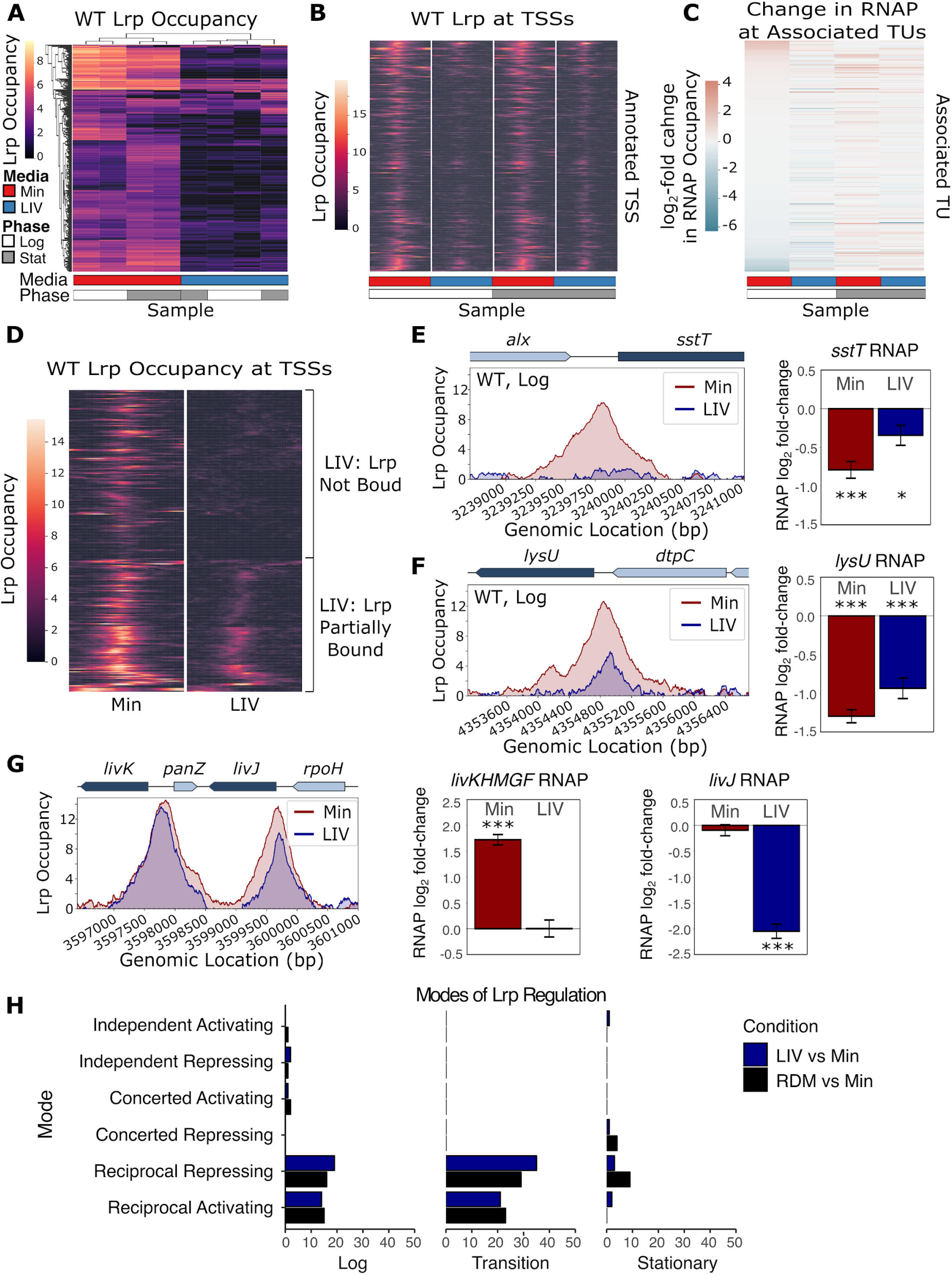
Leucine abrogates or reduces WT Lrp binding to DNA. **A**. Heatmap of WT Lrp occupancy at all significant WT Lrp binding sites across conditions. The y-axis contains all 440 significant WT Lrp binding sites, the x-axis shows the average of both biological replicates for each lineage replicate in each media (Min in red and LIV in blue) and each time-point (mid-log in white and stationary phase in gray), and the color at each location represents the *lrp::scar*-subtracted Lrp occupancy. Both the x- and y-axes are clustered by similarity. **B**. Transcription Start Site (TSS) pile-up plot for each WT Lrp condition. TSS-pile-up plots, where each row represents a Lrp-regulated transcription start site (TSS) and each column is centered on the annotated TSS, show the WT Lrp occupancy within +/- 1kb of that TSS (color), with transcription moving to the right. For each of the four conditions (columns), the WT Lrp occupancy shown represents the mean of both biological replicates of both lineage replicates. **C**. Heatmap of changes in RNAP occupancy, showing the log_2_-fold-change of *lrp::scar*-subtracted RNAP signal relative to input for the transcriptional unit (TU) downstream of the annotated TSS in the corresponding row of the pile-up plot from B), where a positive value (red) indicates activation and a negative value (blue) indicates repression. The color represents the mean of the change in RNAP occupancy across the entire TU for both biological replicates of both lineage replicates. **D**. TSS pile-up plot for WT Lrp in mid-log phase (equivalent to the first two columns of panel B), sorted by the effect of LIV on Lrp binding at each TSS. The data fall into two distinct classes of WT Lrp binding at a TSS: leucine fully abrogates WT Lrp binding (top) or leucine reduces WT Lrp binding to the TSS (bottom). **E**. Example of LIV abrogating WT Lrp binding to a TSS. Left, WT Lrp occupancy at the *sstT* locus at mid-log phase in Min (red) or LIV (blue) media. Right, log_2_-fold-change in RNAP occupancy across the *sstT* TU at mid-log phase in Min (red) or LIV (blue) media, where positive values represent gene activation and negative values represent repression. Error bars represent the standard error of the mean. *=IHW-q-value <0.05, **=IHW-q-value <0.01, and ***IHW-q-value<0.001. **F**. Example of LIV significantly reducing WT Lrp binding to a TSS. Same as E), except at the *lysU* locus. **G**. Example of LIV having minimal effect on WT. Same as E), except at the *livKHMGF-livJ* locus. **H**. Classification of TUs into the six modes of gene regulation based on RNA expression changes and Lrp binding to the promoter regions. Paired Lrp-ChIP-seq and RNA-seq data from Kroner et al. [5] was classified into one of the six categories of Lrp gene regulation (Independent, Concerted, or Reciprocal with Lrp either activating or repressing) based on whether Lrp was bound to the associated TSS and the changes in gene expression were significant. Because that study used Min, LIV, and rich-defined medium (RDM), all genes were classified into the six classes of Lrp regulation by comparing either LIV to Min (blue) or RDM to Min (black). “Transition” indicates the time-point when cells are transitioning from logarithmic growth into stationary phase growth.

We noticed that the TSSs with the highest WT Lrp occupancy in LIV were those with the greatest Lrp-mediated changes in expression at the associated TU (see top and bottom rows of the LIV-Log column in Figure 3B). To further investigate this effect of exogenous leucine on Lrp binding to DNA, we next clustered the WT Lrp-regulated TSSs by similarity in Lrp occupancy profile during exponential growth. These TSSs fell into two distinct classes: those that completely lost Lrp binding in LIV relative to Min (Fig. 3D, top) and those that were partially bound in LIV relative to Min (Fig. 3D, bottom). Figure 3E illustrates WT Lrp occupancy at the *sstT* locus, which exemplifies the class where Lrp binding is abrogated in LIV. Relief of Lrp binding to the *sstT* promoter in LIV also caused reduced repression of *sstT*. In contrast, Figure 3F shows WT Lrp occupancy at the *lysU* locus, where Lrp occupancy is reduced, but not completely abrogated in LIV relative to Min and here the change in *lysU* expression between Min and LIV is not as strong as at *sstT*. Out of all 142 WT Lrp-regulated TUs, there was only one exception to the trend that leucine inhibits Lrp binding to DNA: *livK*-*livJ* (Fig. 3G), which maintained high Lrp occupancy in LIV, albeit still slightly reduced relative to Min. Intriguingly, despite *livKHMGF* and *livJ* having similar Lrp occupancy profiles at their TSSs in Min and LIV, the expression levels are significantly different between these two conditions, indicating that other factors are interacting at these promoters with Lrp, which has also been observed for the *pap* promoter in uropathogenic *E. coli* [20].

Collectively, our findings suggest that the role of leucine in Lrp-mediated gene regulation is to inhibit Lrp binding to DNA (either partially or fully, depending on the site). Such a mechanism would directly contradict the long-standing hypothesis in the field that Lrp utilizes six modes of gene regulation where Lrp can activate or repress expression, and leucine augments Lrp’s effect (concerted), inhibits Lrp (reciprocal), or has no effect on Lrp (independent). To address the discrepancy in a relatively unbiased manner, we turned to the paired Lrp-ChIP-seq and RNA-seq dataset from Kroner *et al* [5] and classified all Direct Lrp-regulated TUs (defined as those with Lrp bound at the promoter that also had significant Lrp-dependent changes in expression in at least one condition) into one of the six modes of regulation (Fig. 3H). The only categories containing substantial numbers of TUs were the Reciprocal Activating and Reciprocal Repressing, further demonstrating that leucine’s role in Lrp gene regulation is almost exclusively to inhibit Lrp, and that any instances of the other four modes of regulation represent highly unusual special cases (and may well arise partly due to indirect effects of Lrp).

### L136R is insensitive to exogenous leucine

Clustering of similar Lrp occupancy patterns in Figure 2B revealed that L136R in LIV media behaved almost identically to WT Lrp in Min media (See Groups III and IV), suggesting that L136R is insensitive to LIV. We further characterized the L136R octamer-only mutant by generating a heatmap of L136R occupancy at all 517 regions for which a significant L136R peak was called in at least one condition (Fig. 4A, right) and compared these findings with the occupancy profile of WT Lrp (Fig. 4A, left). Consistent with our findings from Figure 2B, the L136R mutant binds to DNA with similar intensities in Min and LIV media, unlike WT Lrp, as quantified by pairwise Spearman correlations of Lrp occupancy profiles between conditions in Figure 4B. To investigate whether this leucine-insensitive effect remains true for L136R-mediated gene regulation, we plotted L136R occupancy at all 156 TSSs overlapped by a L136R Lrp peak (Fig. 4C), along with the changes in RNAP occupancy within the associated TU (Fig. 4D). Both the TSS pile-up plots and the changes in RNAP occupancy were similar when comparing Min-Log with LIV-Log, with L136R occupancy decreasing only slightly in LIV-Log relative to Min-Log, demonstrating that L136R is leucine insensitive with respect to both DNA binding and gene regulation.

**Figure 4:**
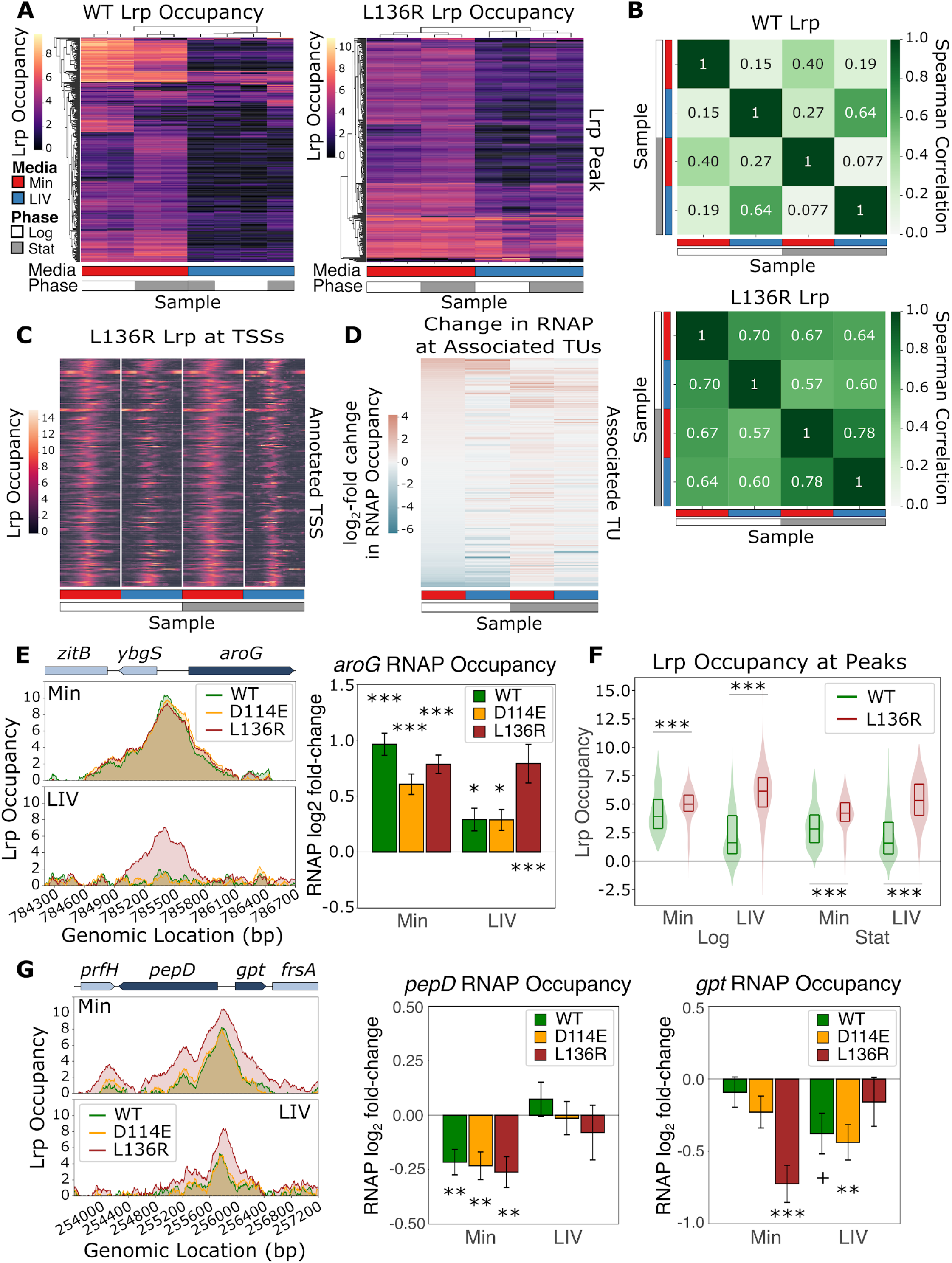
L136R binds more strongly to WT Lrp DNA binding sites, regardless of LIV. **A**. Left, heatmap of WT Lrp (same as Fig. 3A). Right, heatmap of L136R Lrp. The y-axis (right) contains all 517 significant L136R Lrp binding sites in at least one L136R Lrp condition, the x-axis shows the average of both biological replicates for each lineage replicate in each media (Min in red and LIV in blue) and each time-point (mid-log in white and stationary phase in gray), and the color at each location represents the *lrp::scar*-subtracted Lrp occupancy. Both the x- and y-axes are clustered by similarity. **B**. Pairwise Spearman correlations of the WT and L136R heatmaps from A). **C**. TSS-pile-up plots, where each row represents a Lrp-regulated transcription start site (TSS) and each column is centered on the annotated TSS, showing the L136R Lrp occupancy within +/- 1kb of that TSS (color), with transcription moving to the right. For each of the four conditions (columns), the L136R Lrp occupancy (color) shown represents the mean of both biological replicates of both lineage replicates. **D**. Heatmap of changes in RNAP occupancy for each L136R Lrp condition shown at the TUs corresponding to the TSSs in panel C. **E**. Left, Lrp occupancy at mid-log phase in Min (top) and LIV (bottom) at the *aroG* region. Right, changes in RNAP occupancy for the *aroG* TU for each of the three Lrp variants in Min and LIV at mid-log phase. Error bars represent the standard error of the mean. +=IHW-q-value <0.1, *=IHW-q-value <0.05, **=IHW-q-value <0.01, and ***IHW-q-value<0.001. **F**. The violin plots display the overall distribution of Lrp occupancy (WT in green and L136R in red) under each condition, whereas the inner boxplot shows the 25th, median, and 75h percentile of the data. Significance was calculated using the paired Wilcoxon signed-rank test; *** indicates p<0.001. **G**. Same as E), except at the *pepD-gpt* region.

Two interesting, yet related, trends arise in the L136R mutant: in LIV media, L136R Lrp behaves almost identically to WT in Min media, but in Min media, L136R Lrp binds more strongly to canonical Lrp sites than WT Lrp. For example, Figure 4E demonstrates that L136R Lrp in LIV behaves like WT Lrp in Min at the *aroG* region. Here, L136R Lrp occupancy in LIV (bottom) very closely mimics the WT Lrp occupancy in Min media (top). Furthermore, these similar Lrp occupancy traces result in similar effects on gene expression; L136R in LIV media activates expression of *aroG* to a similar extent as WT Lrp in Min media (Fig. 4E). In contrast, WT and D114E Lrp no longer bind to the *aroG* TSS in LIV and also no longer activate expression of *aroG* in LIV media (Fig. 4E). Intriguingly, in Min media, L136R Lrp also binds more strongly to DNA than WT Lrp (Fig. 4F). The *pepD-gpt* region as shown in Figure 4G serves as an example of this trend, with L136R Lrp showing stronger occupancy signal at the *pepD-gpt* region in Min media relative to WT Lrp (top). While the increased occupancy of L136R at the *pepD-gpt* promoters did not have a significant differential effect on *pepD* expression in Min media, it caused a significant increase in *gpt* repression relative to WT Lrp, suggesting that this increased binding to DNA is functional and leads to changes Lrp-mediated gene regulation.

Overall, L136R is leucine insensitive; it binds to existing Lrp sites more strongly in Min media than WT Lrp, and in LIV, L136R behaves almost identically to WT Lrp in Min media. Indeed, L136R-Min had some of the highest Lrp occupancies across all strains and conditions tested (Fig. 2B), despite having a lower Lrp protein level than WT in all conditions tested (Fig. 1C). This suggests that L136R is insensitive to leucine and thus can bind more strongly to its binding sites than WT Lrp, which is inhibited by leucine. It is important to note that, while the Lrp occupancy is higher in L136R samples than WT samples, there do not appear to be many novel DNA binding sites in L136R relative to WT Lrp (Fig. 2C).

### D114E has novel binding sites in the genome and generates additional peaks near existing ones

In contrast to L136R, the D114E “octamer-only” Lrp mutant binds to many novel sites in the genome relative to WT Lrp, in addition to canonical Lrp sites (Fig. 2B: Classes 2, 3, and 7). A heatmap of Lrp occupancy for D114E samples at all 556 regions bound by D114E in at least one condition (compared to only 440 sites for WT Lrp) reveals that exogenous leucine strongly reduces D114E occupancy at its target sites relative to Min media, suggesting that, unlike the L136R octamer-only mutant, the D114E octamer-only mutant is responsive to leucine (Fig. 5A). Similar to WT Lrp, D114E Lrp occupancy at the 160 TSSs overlapped by a D114E peak was reduced in LIV-Log relative to Min-Log, and this reduction in Lrp binding was often accompanied by a reduction in the degree to which RNAP occupancy at the associated TU changed. In general, D114E appears to bind novel sites on the genome in Min media and also generates novel secondary peaks in close proximity to canonical WT Lrp binding sites in Min media, both of which are typically disrupted in LIV media.

**Figure 5:**
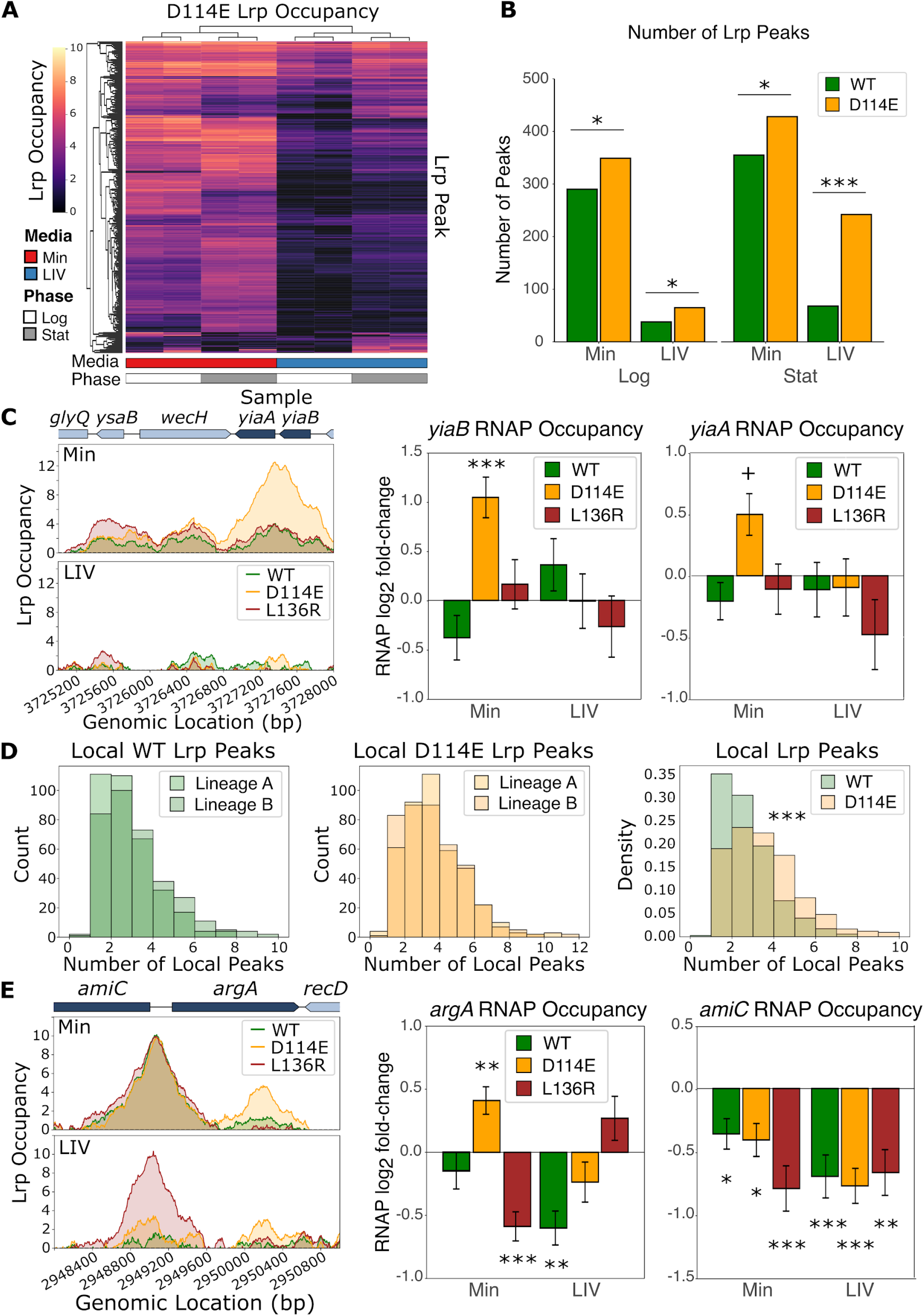
D114E has novel binding sites in the genome and generates additional peaks near existing ones. **A**. Heatmap of D114E Lrp occupancy at all significant D114E Lrp binding sites across conditions, where the y-axis contains all 556 significant D114E Lrp binding sites in at least one D114E Lrp condition, the x-axis shows the average of both biological replicates for each lineage replicate in each media (Min in red and LIV in blue) and each time-point (mid-log in white and stationary phase in gray), and the color at each location represents the *lrp::scar*-subtracted Lrp occupancy. Both the x- and y-axes are clustered by similarity. **B**. Total number of peaks in each of the four conditions tested for WT and D114E Lrp. **C**. Left, Lrp occupancy at mid-log phase in Min (top) and LIV (bottom) at the *yiaB-yiaA* region. Changes in RNAP occupancy for the *yiaB* TU (middle) and *yiaA* TU (right) for each of the three Lrp variants in Min and LIV at mid-log phase. **D**. Histograms of local peak counts for WT lineage replicates (left) and D114E lineage replicates (middle) are shown; see Materials and Methods for details. Right, histogram of the density of local peak counts for WT vs D114E, which show a significant difference in mean counts (p<0.001, permutation test). **E**. Same as C), except at the *amiC-argA* region.

Figure 5B demonstrates that D114E Lrp generates additional Lrp sites relative to WT Lrp in every condition tested. Across all conditions, D114E had 208 unique peaks relative to WT Lrp, but only 35 of these peaks (16.8%) overlapped with an annotated TSS. An example of one such novel D114E Lrp peak is shown in Figure 5C, where additional binding forms in Min media at the *yiaB-yiaA* region. Unlike WT and L136R Lrp, D114E Lrp forms a strong peak at the *yiaB* and *yiaA* promoters, which is also accompanied by strong activation of both *yiaB* and *yiaA* in Min media, while WT and L136R do not significantly alter expression. These unique D114E sites may have arisen in part due to higher Lrp protein levels in the D114E strains than WT or L136R Lrp strains (Fig. 1C).

We also observed that D114E Lrp often formed novel local secondary peaks in addition to binding to the canonical WT Lrp sites. For each lineage replicate (A and B) of both WT and D114E Lrp strains, we calculated the distribution of the number of local Lrp peaks at each defined Lrp peak and found that WT Lrp predominantly has only one or two local peaks, whereas D114E is prone to having 1-4 local Lrp peaks (Fig. 5D).As an example of D114E forming an extra secondary peak near a canonical Lrp binding site, Figure 5E shows that D114E forms the canonical Lrp peak at the *argA* promoter (along with WT and L136R) in Min media, but also forms an additional peak within the *argA* ORF, where neither WT nor L136R Lrp are bound. Intriguingly, this additional D114E peak at *argA* is associated with significant Lrp-mediated activation of argA, whereas WT and L136R, which lack this additional peak, both repress *argA* in Min media (Fig. 5E). While the exact molecular mechanism giving rise to the extra binding sites in D114E Lrp strains is unclear, the most obvious hypothesis is that the multiple local peaks represent higher-order (larger than octamer) Lrp oligomers which can then bridge multiple sites on the genome over kilobase length scales. Thus, while D114E was originally characterized as an “octamer-only” Lrp mutant, the D114E mutation actually most likely stabilizes higher-order Lrp oligomers in Min media relative to WT Lrp, allowing it to occupy these additional sites next to canonical Lrp peaks.

### Lrp forms higher-order oligomers and bridges DNA by binding to one or more nearby regions

While D114E exhibits a propensity for forming secondary peaks within close proximity to primary Lrp peaks on TSSs, WT Lrp also forms both double peaks (Fig. 6A) and secondary peaks (Fig. 6B) in Min media, but only at a subset of sites (Fig. 5D). Interestingly, the midpoints of the double and secondary Lrp peaks are almost always separated by 1500bp (e.g., Figs. 6A and 6B). One particularly striking example is the *fadR-ycgB-dadAX* region, which contains a strong primary Lrp peak (Peak 3) at the divergent *ycgB* and *dadAX* promoters and three secondary peaks (two upstream and one downstream) in Min media (Fig. 6C, bottom), separated by 1200-1800 bp. We chose to study the fadR-ycgB-*dadAX* region in order to understand what role, if any, the observed local secondary peaks play on Lrp binding within the primary Lrp peak and vice versa If the Lrp peaks show correlated changes in binding across perturbations of different sites, it would support our hypothesis that Lrp (particularly D114E) forms higher-order oligomers along the DNA, especially in Min media, where binding to DNA is generally stronger. We thus constructed four strains (WT, Lrp5 scrambled, Peak 3 scrambled, and all secondary peaks scrambled) (Fig. 6C, top) with permutations to different portions of the fadR-ycgB-*dadAX* region, and performed Lrp-ChIP-qPCR on two separately-constructed lineage replicates of each strain to measure changes in Lrp occupancy in Min-Log relative to two control regions (*cysG* and *mdoG*). Within the primary peak (Peak 3), there are 11 previously characterized Lrp dimer binding sites, and the 17bp Lrp5 binding site is essential for de-represion of *dadAX* in LIV, but is typically bound by Lrp in both Min and LIV media [21]. Because Peak 3 is about 1kb wide and sits on the divergent *ycgB* and *dadAX* promoters, we scrambled the regions not overlapping the -35 and -10 sites and made synonymous mutations in the open reading frames (ORFs) so as to minimally disrupt the proteins synthesized from these genes. Similarly, because the secondary peaks were each about 500bp and overlapped *fadR, ycgB*, and *dadX* ORFs, we made synonymous mutations such that the encoded proteins would be intact but that the Lrp binding sites on the DNA would be disrupted.

**Figure 6:**
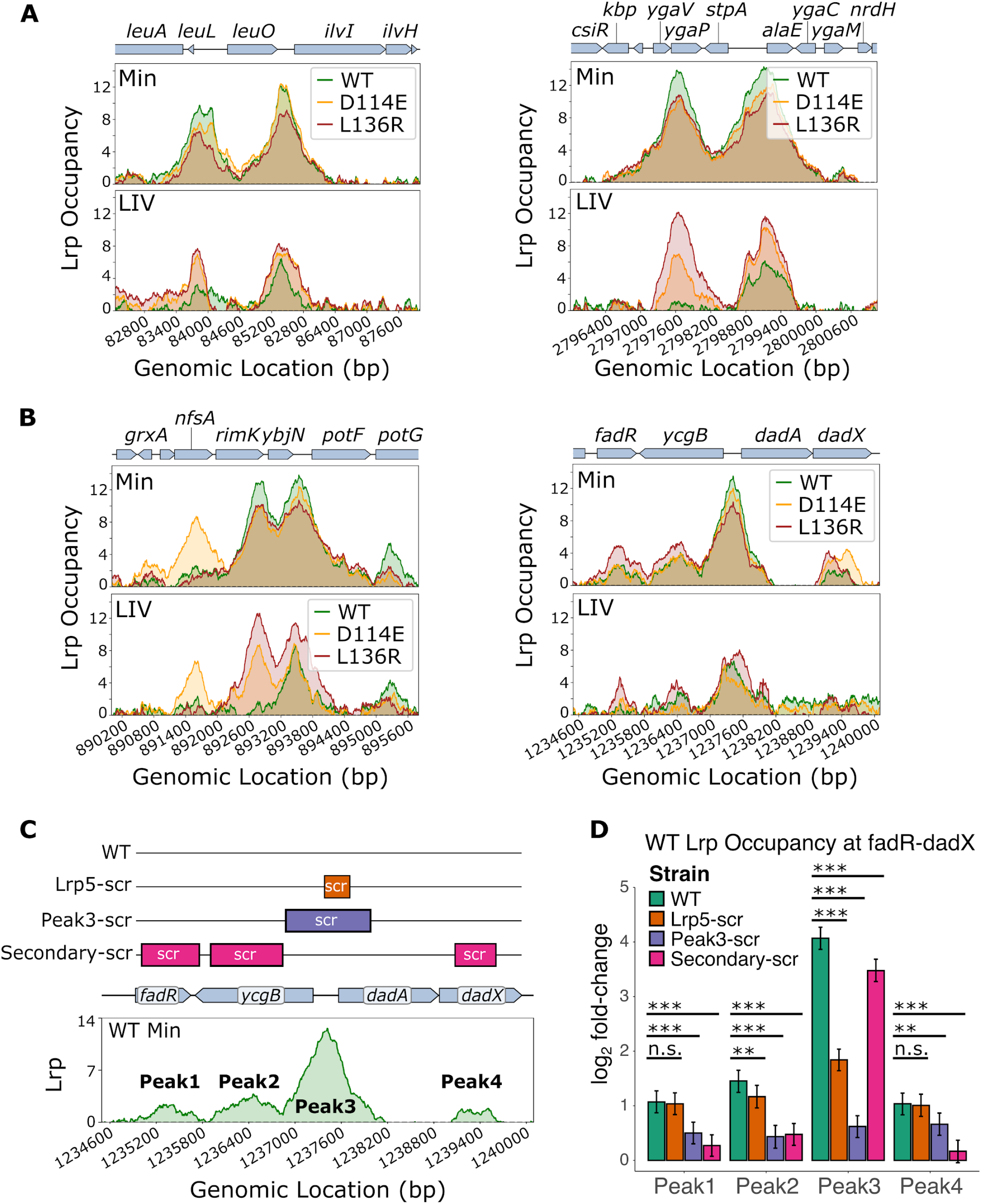
Lrp bridges the DNA locally by binding to one or more nearby regions. **A**. Examples of double Lrp peaks, showing Lrp occupancy at mid-log phase in Min (top) and LIV (bottom) at the *ilvIH* region (left) and *stpA-alaE* region (right). **B**. Examples of Lrp secondary peaks. Same as A, except for the *potF* region (left) and *dadAX* region (right). **C**. Top, diagram of strain mutations. All constructs have a FRT-flanked *KanR* marker immediately downstream of dadX. Lrp5-scr scrambles the 17bp Lrp5 binding site within Peak3. Peak3-scrambled was carefully constructed so as to keep the *ycgB* and *dadAX* promoter elements and ribosome binding sites intact. All Peak3-scramble and Secondary-scramble sequences that overlapped an open reading frame were synonymous mutations such that the amino acid sequences remained intact, but the DNA sequences varied to prevent or reduce Lrp binding. Bottom, WT Lrp occupancy in minimal media at mid-log phase at the *fadR-ycgB-dadAX* region with all four peaks labeled accordingly. **D**. Lrp ChIP-qPCR showing occupancy at each of the four WT Lrp binding sites within the fadR-ycgB-dadAX region in the presence of the sequence variations shown in panel C. Bars shown are the means from a Bayesian model of the log_2_-fold-change of Lrp-ChIP-qPCR at Peaks 1-4 relative to Input, as compared to the average of two control regions where Lrp is known to not bind; error bars show a 95% credible interval for each case.

We observed that scrambling just the 17bp Lrp5 site in the *ycgB-dadA* intergenic region significantly reduced Lrp binding to Peak 3, while not significantly affecting Lrp binding to the secondary peaks (except Peak 2) (Fig. 6D). Furthermore, scrambling the Peak 3 sequence almost completely abrogated Lrp binding to Peak 3 (as expected), but also significantly reduced Lrp binding at the secondary peaks, suggesting that Lrp accumulation at the main peak in the divergent ycgB-*dadAX* regulatory region is required for secondary Lrp peak formation. Interestingly, scrambling all of the secondary peaks also significantly reduced Lrp binding to the primary peak (Fig. 6D). Since perturbing Lrp accumulation at one peak affects Lrp binding at the others nearby, it is highly likely that Lrp at the primary and secondary Lrp peaks makes up a higher-order Lrp oligomer that bridges these DNA sites together in 3-D space. Furthermore, scrambling of Lrp5 was sufficient to greatly reduce occupancy at the primary Lrp peak, suggesting that the Lrp5 site (17bp) serves as the seed sequence that nucleates Lrp binding to the primary peak (∼1kb). While *E. coli* Lrp has only previously been shown to exist in oligomers as large as a hexadecamer, these findings may suggest that *E. coli* Lrp can potentially form even higher-order oligomeric complexes along specific stretches of DNA, or that at sites such as the fadR-ycgB-*dadAX* region Lrp exhibits heterogeneous long-range contact formation across a population of cells. These findings also strengthen our conclusion that D114E, which forms novel secondary peaks near many canonical Lrp binding regions, stabilizes the higher-order Lrp oligomeric state that bridges DNA in Min media.

## Discussion

*E. coli* Lrp has long been hypothesized to be in an equilibrium between hexadecamers (16mers) and octamers (8mer), with exogenous leucine strongly shifting the equilibrium towards the octameric state. However, these findings were insufficient to mechanistically explain how Lrp utilizes six different modes of regulation of its target genes, where Lrp activates or represses and leucine has no effect, augments Lrp activity, or inhibits Lrp activity. Through careful construction of strains harboring both the native *lrp* promoter and gene body at a safe genetic locus in a background with a fully intact branched-chain amino acid metabolism, we were able to reproduce old findings, while also providing insight into the mechanism of Lrp regulation of its target promoters. We show that the role of exogenous leucine is to inhibit WT Lrp binding to DNA, and demonstrate both on our dataset and a previously published dataset that WT Lrp only actually utilizes two modes of gene regulation at its target promoters: Lrp activates or represses transcription, while exogenous leucine inhibits Lrp activity by reducing its affinity for DNA. The other four modes of Lrp regulation at target promoters reported in the literature likely arose from indirect effects or specialized interactions also involving another regulator at a handful of loci.

By coupling small-scale experiments with high-throughput ChIP-seq on WT Lrp as well as three oligomeric mutants under various conditions, we were also able to connect the effects of leucine, oligomeric state, and DNA binding. The dimer-only Lrp (ΔC11) was effectively unable to bind DNA, despite being expressed at only slightly lower levels than WT Lrp at mid-log phase, corroborating the *in vitro* gel shift results from Chen et al. [12]. Previous studies of D114E and L136R suggested that both mutations locked Lrp in an “octamer-only” state, and analyses of regulation at the *ilvIH* promoter (where the mutants were originally identified) suggested that both mutants also had the same regulatory function [13,15]. Despite using different techniques, our experiments replicated the expected regulatory effects of WT, D114E, and L136R Lrp on *ilvIH* in Min and LIV media, while also demonstrating that these two “octamer-only” mutants actually behave quite differently globally, despite having similar binding and regulatory behaviors at a few specific promoters. L136R appears to be a truly leucine-insensitive Lrp mutant, binding more strongly than WT Lrp to DNA in Min media, and, in LIV, behaving like WT Lrp in Min media. Conversely, D114E responds almost as strongly to exogenous leucine as WT Lrp, but overall binds to more sites in the genome and forms secondary peaks near canonical Lrp sites. These findings suggest that D114E is indeed an oligomeric mutant, but likely not octamer-only; instead we propose that D114E favors the formation of even higher-order oligomers (e.g. hexadecamers) in Min media that bridge DNA. Our ChIP-qPCR experiments on strains with mutated sequences at the locations of secondary Lrp peaks demonstrate that these local peaks likely interact with each other, and provide further evidence for these higher-order Lrp oligomers.

*E. coli* Lrp belongs to an ancient and highly conserved class of proteins called Feast-Famine Response Proteins (FFRPs), with homologs in almost all sequenced archaea and nearly half of all sequenced bacteria [3]. Across all FFRPs, four small molecule effector binding sites have been identified [22]. Two such sites (Type I and Type III) are found in *E. coli* Lrp, where the Type I site is the highly conserved high-affinity effector binding site and Type III is less conserved and serves as a low-affinity effector binding site [6]. Previous studies have reported that residue D114 in Lrp is part of the low-affinity site and contributes to higher-order Lrp oligomerization [13]. Thus, while our results show that D114E is not an octamer-only mutant, it makes sense that this mutation stabilizes higher-order oligomers (e.g. hexadecamers). Conversely, the Lrp L136R mutation is in a highly conserved region that aligns with Type I (high affinity) effector binding sites in other FFRP homologs [22–24], which is consistent with our findings that L136R Lrp is leucine-insensitive. Collectively, our findings demonstrate that neither D114E nor L136R are octamer-only mutants, but rather, that their unique phenotypes are connected to the roles of high and low affinity effector binding sites on Lrp in altering DNA binding (high affinity site, L136R) and oligomerization (low affinity site, D114E). We expect that future experiments more directly probing the oligomeric state of Lrp at different loci under different conditions, and directly identifying any long-range contacts formed by Lrp oligomers, will provide further mechanistic insight into gene regulation by Lrp and other FFRPs. In addition, as the investigation provided here was still performed under artificial conditions (defined media with/without branched chain amino acid supplementation), it remains to be seen how nutrient conditions more commonly encountered in *E. coli*’s evolved ecological niche would be detected by Lrp and alter Lrp-dependent regulation.

## Materials and Methods

### Bacterial Strains and Culturing

The parental strain, RL3000 lrp::scar was generated via P1vir transduction of *lrp::KanR* from the Keio collection [25] strain JW0872 into RL3000 followed by electroporation of pCP20 to flip out the FRT-flanked KanR marker. From here, *thyA::KanR* was transduced from the Keio collection strain JW2795 via P1vir transduction [26] and selected on LB plates supplemented with kanamycin and 20 μg/mL thymidine, followed by electroporation of pKD46 [27]. Plasmids containing *lrp* (with its native promoter) flanked by strong bidirectional terminators and linked to *thyA* were generated for WT *lrp* and each of the *lrp* mutants or *thyA* only for the Δ*lrp* strains. Final strains were constructed via lambda red recombination of the *thyA-lrp* cassette into the pKD46-bearing lrp::scar, thyA::KanR strains, selected on LB lacking thymidine, and confirmed via colony PCR and Sanger sequencing. RL3000, *lacZ::CAT, p*_*oppA*_*-lacZ*, and RL3000, *lacZ::CAT, p*_*alaE*_*-lacZ* strains used for the Miller Assays were generated by first using lambda red recombination to replace native *lacZ* with *CAT* in the RL3000 lrp::scar parent, followed by introduction of thyA::*kanR* via P1vir transduction. From here, a cassette containing *lacZ* linked to loxP-flanked *specR* was recombined into the genome at either native p_*oppA*_ or p_*alaE*_, such that the cassette was integrated between the promoter and translation start codon of *oppA* or *alaE*. Finally, *specR* was flipped out via induction of Cre recombinase from an arabinose-inducible plasmid, and the appropriate *lrp* mutant was integrated into the *thyA* locus using P1vir from the *thyA-lrp* strains from above. All cloning was performed in standard LB media supplemented with appropriate antibiotics. All strains used in experimental measurements (with the exception of the parental RL3000) were constructed in duplicate to ensure the robustness of our results; we refer to the independent but nominally isogenic variants of each genotype as separate lineages.

For physiological experiments such as ChIP-seq, single colonies were inoculated into 0.04% glycerol-ammonia M9 minimal media and grown overnight at 37°C. The next day, these overnight cultures were diluted 1:200 into 0.4%-glycerol-ammonia M9 media (Min) or 0.4%-glycerol-ammonia M9 supplemented with 0.2% each of L-leucine, L-valine, and L-isoleucine (LIV). Log-phase samples were collected at OD_600_=0.2 and stationary-phase samples were collected approximately 12 hours after the log-phase time-point. Each “biological replicate” for such experiments refers to an experiment performed on a separate day arising from a separate colony grown as described above.

### Miller Assays

Single colonies from LB plates were inoculated into 3mL 0.04%-glycerol-ammonia M9 minimal media and grown at 37°C overnight. The next day, 5μL cells were added to 1mL of Min or LIV minimal media (see above) in 2mL wells of a deep 96-well plate and grown at 37°C with shaking until approximately the desired OD_600_ was reached (around 0.2). At this point, 80μL cells were taken and added to a clear 96-well plate and the OD_600_ of each well was measured on a plate reader. 120μL of the **β**-Galactosidase master mix (80μL Z-Buffer with 2.7μL/mL **β**-mercaptoethanol, 30μL Z buffer with 4mg/mL ortho-nitrophenol-galactoside (ONPG), 8μL PopCulture detergent (EMD Millipore Corp., Billerica, MA), and 2μL10mg/mL lysozyme in water, where Z-Buffer consists of 60mM Na_2_HPO_4_, 40mM NaH_2_PO_4_, 10mM KCl, and 1mM MgSO_4_) was then carefully mixed into each well with cells and the OD_550_ and OD_420_ were monitored every two minutes for one hour in a plate reader with shaking between readings. RL3000 in the presence of 1mM IPTG was used as a positive control, whereas RL3000, *lacZ::CAT* was used as the negative control in these experiments. This protocol was adapted from [28] and [29].

### Western Blots

Single colonies from each strain were inoculated into 5mL LB and grown overnight at 37°C, back-diluted 1:200 into 3mL 0.04% Glycerol-Ammonia M9 media and grown again overnight at 37°C. The OD_600_ of the resulting overnight cultures was measured and the appropriate amount of cells were added each into 5mL 0.4% Glycerol-Ammonia M9 media (Min) and 5mL 0.4% Glycerol-Ammonia M9 media supplemented with 0.2% each of L-leucine, L-isoleucine, and L-valine (LIV) such that the starting OD_600_ in these cultures was 0.003. Cultures were grown at 37°C with shaking until mid-log phase (OD_600_ = 0.2), at which point 2mL was spun down and cell pellets were flash-frozen at -20°C. Approximately 12 hours after the mid-log time-point was taken, the OD_600_ was again measured, 1mL of these stationary phase samples were spun down, and cell pellets were flash-frozen at -20°C.

Cell pellets were resuspended in 1x Laemmli Loading Buffer (5µL 4x Laemmli Buffer: 1µL 1M DTT: 14µL 1xPBS) such that the concentration of cells was OD_600_=10/mL. Solutions of 0.5µM, 1µM, and 5µM solutions of purified native Lrp in 1x Laemmli Loading Buffer in order to calculate the amounts of Lrp in each sample and across blots. Samples were denatured at 99°C for 10min. 7µL of Bio-rad Precision-Plus Protein All-Blue Standards and 10µL each sample were then loaded into the wells of 15well 4-20% Bio-rad Mini-PROTEAN TGX Stain-Free Gels and electrophoresed at 175V for 45min at room temperature in Tris-Glycine-SDS Running Buffer. Gels were then rinsed in milliQ water and total protein was imaged with the 5min Stain-Free gel protocol on the Bio-Rad ChemiDoc MP Imaging System. Proteins were transferred onto Bio-rad Immuno-Blot PVDF Membranes (pre-activated in 100% methanol) via the wet transfer technique in Tris-Glycine-Methanol Transfer Buffer at 60V for one hour at 4°C. PVDF membranes were briefly rinsed in milliQ water and then incubated with 30mL 3% non-fat milk in TBST (Tris-Buffered Saline with Tween20) with gentle staking at room temperature for 30 minutes. Excess milk was removed from the membrane and 10mL 1:4,000 monoclonal Lrp antibody in 3% non-fat milk in TBST was added to each membrane, incubating for 2 hours at room temperature. Primary antibody solution was removed and 10mL 1:5,000 HRP-goat, anti-mouse antibody in 3% non-fat milk in TBST was added to the membrane and allowed to incubate overnight at 4°C. The next day, PVDF membranes were washed 3x with TBST, allowing the membranes to gently shake at room temperature for 10min between washes. After washing, the PVDF membranes were rinsed briefly with milliQ water and 1mL of MilliPore Immobilon Western Chemiluminescent HRP Substrate was added to the top of the membrane for 1 minute prior to imaging. Images were taken with a two-second exposure in the Chemiluminescence protocol on the Bio-Rad ChemiDoc MP Imaging System.

Raw image (tiff) files from the Lrp western blot images were loaded into ImageJ software and rectangles of the same size were placed over each Lrp band across the blot. Background signal was subtracted and the total pixel density was calculated for each band. Total protein signal was measured in a similar manner by loading the total protein gel image tiff files into ImageJ and, instead of placing a rectangle around a specific band, a rectangle was placed around each lane of the gel. After background subtraction, the total pixel density from each lane was calculated. To normalize Lrp levels and generate relative Lrp abundances on the Western Blots, the Lrp western blot band pixel density was divided by the total protein blot lane pixel density for each sample.

The relative Lrp abundances were then analyzed using a Bayesian model, in which we assumed population-level (fixed) effects for the biological condition (media/growth phase combination), sample genotype, genotype:media interaction, genotype:growth phase interaction, and genotype:condition interaction, as well as group level (random) effects for each specific gel and for the effects of the condition given the strain identity. Data were log_2_-scaled and z-scored prior to model fitting, and then we report the back-transformed parameters in the original data units. The models were fitted using the R package brms [30,31] with 5000 iterations in each of four Monte Carlo chains; convergence was assessed based on the Rhat criterion plus visual inspection of the posterior predictive distributions. We used a normal(0,3) prior for the population-level effects, a normal(0,2) prior for the group-level effects, an lkj(10) prior for correlation parameters, and default priors for all other parameters.

### Paired Lrp-ChIP-seq and RNAP-ChIP-seq

Single colonies of each lineage of each strain were first grown overnight at 37°C in 0.04%Glycerol, 0.4%NH_4_Cl M9 minimal media with shaking, The next day, cells were back-diluted to OD_600_=0.003 into 250mL 0.2%Glycerol, 0.4%NH_4_Cl M9 minimal media (Min) or 250mL 0.2%Glycerol, 0.4%NH_4_Cl M9 minimal media supplemented with 0.2% each of L-leucine, L-isoleucine, and L-valine (LIV) and grown at 37°C with shaking until OD_600_=0.2 (log-phase), at which point 100µL cells were serially diluted in 1x PBS and plated on LB plates in order to calculate CFU/mL for each sample, and three separate aliquots of 30mL cells were added to 810µL 37% formaldehyde solution (final concentration = 1%), and 300µL 1M NaPO_4_, pH=7.4 and incubated with shaking at room temperature for 15 minutes. To quench the crosslinking reactions, 4M Tris, pH=8.0 was added to a final concentration of 280mM and samples were incubated with shaking at room temperature for an additional ten minutes. Cells were spun down at 15,000 x g for 2minutes at 4°C and cell pellets were washed twice with 50mL ice-cold TBS (50mM Tris, pH=7.5, 150mM NaCl). Final cell pellets were resuspended in 1mL TBS, transferred to 1.7mL Eppendorf tubes, and centrifuged for 3 minutes at 12,000 x g at 4°C. Upon removal of supernatant, cells were flash-frozen in a dry-ice ethanol bath and stored at -80°C. The same protocol was followed for the stationary phase time-point, which occurred 12hrs after removal of the log-phase aliquots. Note that for each sample and time-point, three cell pellets were generated.

Two of the frozen cell pellets for each log-phase sample were resuspended and combined in a total of 600µL Lysis Buffer and vortexed for 3 seconds. For stationary-phase samples, one pellet was resuspended in 1mL PBS and 100µL of this solution was added to 500µL Lysis Buffer. All samples were incubated at 37°C for 30min and then placed on ice. Samples were kept cold and sonicated for three bursts of 10s ON, 10s OFF at 25% power. 6µL RNaseA (10mg/mL), 5.4µL 100mM MnCl _2_, 4.5µL 100mM CaCl_2_, and 6µL DNaseI were added to each sample and mixed gently. Samples were incubated on ice for 15min and quenched with 50µL 0.5M EDTA, pH=8.0. Cell debris were removed via centrifugation at 16,000 x g for 10min at 4°C.

Cell lysates were added to 50µL pre-washed Protein G beads (supernatant from 50µL beads was discarded and beads were washed 3x with 1mL PBS+0.1% Tween20, final beads were resuspended with wide-bore tips in 50µL PBS+0.1% Tween20) and gently rocked at room temperature for one hour to pre-clear the lysates. Samples were then placed on a magnetic stand and the supernatant was transferred to a fresh tube. 50µL pre-cleared lysate was set aside as the Input sample and mixed with 450µL Elution Buffer. Input samples were incubated for 12-16hrs overnight at 65°C to reverse crosslinks. The remainder of the cell lysate was split in half: 275µL for the RNAP-ChIP and 275µL for the Lrp-ChIP.

For the Lrp-ChIP, 100µL Protein G beads were washed 3x in 1mL PBS+0.1%Tween20 and resuspended in 100µL PBS+0.1%Tween20. These pre-washed beads were incubated with 10µg Lrp monoclonal antibody for 10 minutes at room temperature with gentle shaking. The 275µL Lrp-ChIP sample was then added to the bead-antibody mixture and incubated at room temperature for 1 hour with gentle rocking.

The beads were then washed 3x with 1mL PBS+0.1%Tween20. 500µL Elution Buffer was added to the beads and samples were incubated at 65°C for 25 minutes with vortexing every 5 minutes. Samples were placed on a magnet stand and supernatant was placed into a fresh tube. Crosslinks were reversed by incubating overnight (12-16hrs) at 65°C.

The 275µL of RNAP-ChIP samples were added to 275µL 2x Mooney IP Buffer and incubated with 10µg RNAP-ChIP antibody overnight with gentle rocking at 4°C. The next day, 50µL Protein G beads (per sample) were placed on a magnet stand and washed once with 1mL 1x Mooney IP Buffer and resuspended in 50µL 1x Mooney IP Buffer. These pre-washed beads were then added to the lysate-antibody mixture and incubated for 2hours at 4°C with gentle rocking. For the next wash steps, 1mL of buffer was added, tubes were mixed thoroughly by inversion, placed back on the magnet stand, and supernatant discarded in the following order: IP Wash Buffer A, IP Wash Buffer B, IP Wash Buffer C, 1x Mooney IP Buffer with 1mM EDTA, and 1x TE. Beads were then resuspended in 500µL Elution Buffer and incubated at 65°C for 25minutes with vortexing every 5 minutes. Finally, samples were placed on a magnet srand and supernatant was removed to a fresh tube. Crosslinks were reversed by incubating overnight (12-16hrs) at 65°C.

After treatment with 10µL RNaseA (10mg/mL) for 2hours at 37°C followed by 10µL Proteinase K for 2hours at 50°C. DNA from the Input, Lrp-ChIP, and RNAP-ChIP tubes was then extracted via phenol-chloroform precipitation and concentrated via ethanol precipitation.

### Preparation of DNA samples for Illumina Sequencing

Concentrations of DNA samples were determined using dsDNA Quantifluor (Promega) following manufacturer’s instructions, and samples were diluted to 5ng//µL prior to performing Illumina sequencing preps with the NEBNext Ultra II DNA Library Prep Kit, following manufacturer’s instructions, with the following modification to the first bead-cleanup step: 68µL isopropanol and 174µL magnetic beads were added to each sample and then washed twice with fresh 80% ethanol as normal. All sequencing preps were performed using the Opentrons2 Liquid-handling robot up through the final PCR amplification step. Final bead cleanups were performed by hand, and DNA concentrations were again determined via dsDNA Quantifluor (Promega) following manufacturer’s instructions. Samples were pooled into a 5nM library and sequenced on an Illumina NovaSeq 6000.

### Processing of Illumina reads and peak calling

Read preprocessing, alignment, and initial quantitation was performed using version 2.3.6 of the IPOD-HR [29] analysis pipeline (https://github.com/freddolino-lab/ipod), up to but not including the ChIP subtraction stage. In brief, reads are clipped to remove identifiable Illumina adapter sequences, quality trimmed from their 3’ ends, and then aligned to the E. coli MG1655 genome (version U00096.3) using bowtie2. Coverages were then calculated for each sample across the genome at 5 bp intervals, and the coverages quantile normalized separately for each condition/genotype combination. After quantile normalization, the median value of each sample was set to 100 by rescaling the data, and a pseudocount of 0.25 added to each position. Log_2_ ratios for each non-input sample were calculated relative to the matched input sample; values were then averaged across biological replicates. The averages were subsequently converted to robust z-scores.

For the Lrp ChIP data, to remove any contributions from nonspecific antibody background, the position-wise robust z-scores from *lrp* knockout cells under the same condition were subtracted from the robust z-scores for each sample of interest; the background-subtracted robust z-scores were then used for analysis throughout the text unless otherwise noted.

Lrp peaks were called for each sample for a range of z-score thresholds from 1-8 in intervals of 0.1. We then determined (and subsequently applied) an optimal peak z-score threshold for each sample by maximizing the Kullback-Leibler divergence between the distributions of occupancy scores for peak vs. non-peak regions of the genome (with the occupancy scores discretized into bins at unit increments from -10 to 10, with a pseudocount of 1 added to each bin). Final peak lists for each sample were generated by keeping only the peaks that occurred in both lineages of a single condition with at least a 50% reciprocal overlap (i.e. WT_min_log). Heatmaps of Lrp occupancy at each peak were generated by plotting the average Lrp occupancy across the length of a peak.

### TSS pileup plots

Occupancies around transcription start sites were collected on 2 kb windows centered on all RegulonDB-annotated transcription start sites [32], and then plotted using the clustermap function of seaborn, building on the numpy, scipy, matplotlib, and pandas libraries [33–37].

### RNAP-ChIP analysis

RNAP ChIP reads were processed as described above up through the read alignment stage, and then coverage of transcriptional units (from RegulonDB [32]) calculated using the summarizeOverlaps function of the R package GenomicAlignments [38], using mode “IntersectionStrict”. Differential expression calling and fold change estimation between conditions was then performed using deseq2 [39], with multiple hypothesis testing correction using the IHW method [40].

### Calculation of the distribution of the number of local Lrp peaks

To calculate the number of sub-peaks within each called peak region, we first defined a window around each peak call that contained the full peak plus 2 kb of padding in either direction (using `bedtools slop`), and then smoothed the occupancy with a Hann filter over a 500 bp window (using the `scipy.signal.convolve` function), and flagged peaks using the scipy.signal.find_peaks function with parameters `height=1, distance=50` (corresponding to a 250 bp range between peaks, as data points were taken at 5 bp intervals). The numbers of distinct peaks identified in each window were used to generate the distributions shown in Fig. 5C.

### Lrp-ChIP-qPCR

Two independent lineages (A and B) of *WT fadR-ycgB-dadAX-KanR, Lrp5_scr–fadR-ycgB-dadAX-KanR, Peak3_scr-fadR-ycgB-dadAX-KanR*, and *Secondary_scr-fadR-ycgB-dadAX-KanrR* were generated at the native genomic *fadR-ycgB-dadAX* locus and each strain was grown to mid-log phase in Min media as described in “Preparation of samples for paired Lrp-ChIP-seq and RNAP-ChIP-seq” above. Input DNA samples were diluted 1:150 in 1xTEe such that their concentrations were in the same range as the Lrp-ChIP samples. qPCR was performed on three technical replicates of each sample with Bio-Rad iTaq Universal SYBR Green Supermix according to manufacturer’s instructions and samples were run on the Bio-Rad CFX Opus 384 Real-Time PCR instrument to determine Cq values. Lrp-ChIP Cq values at each locus were offset by the mean Cq value of two control regions: *cysG* and *mdoG*. See Table S3 for primer sequences. To ensure that primer efficiencies were comparable across samples/mutants, we ran qPCR with all appropriate primer pairs on a 10-fold dilution series of the input samples (ranging from 1 to 10^−8^) and analyzed the slopes of the resulting Cq values vs. log_2_ dilutions. Because all of the slopes were highly comparable (−2 +/- 0.1), we did not further normalize Cq values by primer efficiency.

The ChIP-qPCR data were analyzed using a Bayesian model in which the log_2_ fold changes (i.e., difference between ChIP and input Cq values, after correction for the control regions) were treated using a linear model with normally distributed residuals. We included population-level terms for the peak identity (i.e. primer pair), peak:sample interaction (the key parameter of interest), and sample:lineage interaction (accounting for variability across biological replicates), as well as group-level effects for each peak:sample:lineage combination and each peak:technical replicate combination. Models were fit using brms [30,31] using default parameters, and fit quality assessed using the Rhat criterion and visual inspection of the posterior predictive distributions. We used normal(0,10) priors for the peak terms, normal(0,2) priors for the other population-level terms, and brms default priors for all other model terms.

## Acknowledgements

This work was supported by the National Institutes of Health [R35GM128637 to P.L.F.], a National Science Foundation Graduate Research Fellowship [DGE 1256260; to C.A.Z.], an NIH Michigan Predoctoral Training in Genetics Grant [T32GM007544 to C.A.Z.], and a University of Michigan Rackham Predoctoral Fellowship to C.A.Z. We are grateful to Dr. Jeremy Schroeder for coding assistance in developing the version of the IPOD-HR postprocessing pipeline that was used here.

